# The complete mitochondrial genome of Melon thrips, *Thrips palmi* (Thripinae) and comparative analysis: A vector for Tospoviruses

**DOI:** 10.1101/342519

**Authors:** Rajasree Chakraborty, Kaomud Tyagi, Shantanu Kundu, Iftikar Rahaman, Devkant Singha, Kailash Chandra, Srinivas Patnaik, Vikas Kumar

## Abstract

The melon thrips, *Thrips palmi* is a serious pest and vector for plant viruses on a wide range of economically important crops. DNA barcoding evidenced the presence of cryptic diversity in *T. palmi* and that warrants exhaustive molecular data. Our present study is on decoding the first complete mitochondrial genome of *T. palmi* (15,333 bp) through NGS technology. The mitogenome contains 37 genes, including 13 PCGs, two rRNAs, 22 tRNAs, and two control regions. The comparative analyses were conducted for gene arrangements, nucleotide composition, codon usage and phylogenetic relationship with other thrips mitogenomes. The nucleotide composition was 78.29% AT, and 21.72% GC with positive AT skewness (0.09) and negative GC skewness (−0.06). The ATN initiation codons were observed in 12 PCGs except *cox1* with unique start codon (TTG). The RSCU analysis revealed Phe, Leu, Ile, Tyr, Asn, Lys and Met were the most frequently used amino acids in all PCGs. The codon CGG (Arg) was absent in *T. palmi* as compared to other thrips mitogenomes. The Ka/Ks ratio ranges from 0.078 in *cox1* to 0.913 in *atp8*. We observed the typical cloverleaf secondary structure in most of the tRNA genes with a few exceptions; absence of DHU stem and loop in *trnV* and *trnS*, absence of DHU loop in *trnE*, lack of T-arm and loop in *trnN*. The position of *trnS1* (between *cox3* and CR2) is unique in *T. palmi* among all the studied thrips mitogenomes. The mitogenome contained 24 intergenic spacer regions and 12 overlapping regions. The CR2 is 63.77% similar to CR1, indicating a possible duplication and translocation in control region. Both the ML and BI phylogenetic trees revealed the close relationships of *Thrips* with *Scirtothrips* as compared to *Frankliniella*. Thus, more mitogenomes on the diverse thrips species is required to understand the in-depth phylogenetic and evolutionary relationships.

## Introduction

The members of insect order Thysanoptera (commonly known as thrips) are usually tiny, fringe winged and are classified into nine families within two suborders [1]. The family Thripidae is the most diverse family and further divided into four subfamilies (Dendrothripinae, Panchaetothripinae, Sericothripinae and Thripinae). Thrips are widely distributed throughout the world and known by 6154 species, of which 739 species are reported from India [2]. Thrips are one of the major sucking pests and sole transmitters of Tospoviruses (Family Bunyaviridae) on a wide number of agricultural and horticultural crops [3]. Fifteen species are reported as vectors of Tospoviruses, of which six species are known from India (*Ceratothripoides claratris*, *Frankliniella occidentalis, F. schultzei*, *Scirtothrips dorsalis*, *Thrips palmi* and *T. tabaci*) [4-5]. Due to their economic importance, the accurate identification of these species is the basic necessity for the study of their disease transmission efficiency in crops and implementation of the effective management strategies. The morphological identification is time consuming and challenging because of cryptic behaviour and overlapping geographical distributions. Therefore, modern tools such as DNA barcoding have been applied for thrips taxonomy and phylogenetic studies [6–8].

*Thrips palmi* is commonly known as Melon thrips, one of the major pest on agricultural crops and has been reported to vector at least four tospoviruses; Calla lily chlorotic spot virus (CCSV), Groundnut bud necrosis virus (GBNV), Melon yellow spot virus (MYSV), Watermelon silver mottle virus (WSMV) [4]. *Thrips palmi* is widely distributed and highly polyphagous species and is often confused with the *T. flavus* and *T. alatus* [9]. Recent DNA barcoding studies on thrips revealed four MOTUs (Molecular operations Taxonomic Units) in *T. palmi* (*T. palmi* Ia1, *T. palmi* IIa1, *T. palmi* Ib1, and *T. palmi* Ib2) representing multiple cryptic species [8]. Considering these hitches, more molecular data on this species is required to understand the cryptic speciation and evolutionary affiliations.

Mitochondrial genome data have been widely used for phylogenetic, evolutionary studies, and population genetics in insects [10–11]. The circular mitochondrial genome of insects was represented by 37 genes, including 13 PCGs, large and small ribosomal RNA (rrnL and rrnS) genes, 22 transfer RNA (tRNAs) genes and variable number of control regions. However, the availability of mitogenomes in thrips is limited and six mitogenomes of five species (*Anaphothrips obscurus, Frankliniella intonsa, Frankliniella occidentalis, Scirtothrips dorsalis* and *Thrips imaginis*) are available in the GenBank database [12–16].

In the present study, we sequenced and characterized the complete mitochondrial genome of the melon thrips, *T. palmi* under the family Thripidae using the next generation sequencing (NGS) technology and comparative analysed with other thrips mitogenomes. The comparisons were based on genome arrangement, PCGs, tRNAs, rRNAs, nucleotide composition, codon usage, evolutionary rates, etc. Further, to infer the phylogenetic relationships, 13 PCGs of *T. palmi* and other six mitogenomes of five thrips species were analysed using maximum likelihood (ML) and Bayesian inference (BI). This study will provide a better understanding to the comparative mitochondrial genomics of *T. palmi* with the other thysanopterans.

## Materials and methods

### Sample collection, and DNA extraction

The specimens of *T. palmi* were collected from the Odisha State of India from Eggplant (*Solanum melongena*). The specimens were morphologically identified by first author (K.T) with the available taxonomic keys [9], and preserved in absolute ethyl alcohol at −80°C in Centre for DNA Taxonomy, Molecular Systematics Division, Zoological Survey of India, Kolkata. The DNeasy DNA Extraction kit (Qiagen, Valencia, CA) was used to extract the genomic DNA and the concentration was measured on a quantified by Qubit fluorometer (Thermo Fisher Scientific, MA, USA) using a dsDNA high-sensitivity kit with the standard protocol.

### Mitogenome sequencing and assembly

The complete mitochondrial genome sequencing, assembly and annotation were carried out at the Genotypic Technology Pvt. Ltd. Bangalore, India (http://www.genotypic.co.in/). Whole genome sequencing (WGS) library was prepared with Illumina-compatible NEXTflex Rapid DNA sequencing kit (BIOO Scientific, Austin, Texas, U.S.A.). The DNA was sheared using Covaris S2 sonicator (Covaris, Woburn, Massachusetts, USA) to generate approximate fragment size distribution of 200 bp to 600 bp. The fragment size distribution was checked on Agilent TapeStation and subsequently purified using Highprep magnetic beads (Magbio). Purified fragments were end-repaired, adenylated and ligated to Illumina multiplex barcode adaptors. Adapter-ligated DNA was purified using Highprep beads and the resulted fragments were amplified for eight cycles of PCR using Illumina-compatible primers. Final PCR product was purified, followed by library quality control check. Illumina-compatible sequencing library was quantified by Qubit fluorometer (Thermo Fisher Scientific, MA, USA) and its fragment size distribution was analyzed by Agilent 2200 Tapestation and Agilent Bioanalyzer. Libraries with adapter contamination were pooled and cleaned up with Highprep magnetic beads (Magbio) and then sequenced on Nextseq 500 150X2 chemistry. Raw Sequences were trimmed and filtered using NGS Toolkit. The adapter contamination and low-quality reads with base N’s or more than 70% of bases with a quality score <20 also had been removed. The acquired high quality ∼18 million reads were screened out using Burrows-Wheeler Alignment (BWA) tool [17]. Out of 18 million reads, 0.10% (∼1.8 million) of the reads got aligned then assembled with SPAdes 3.9.0 [18], using default parameters considering *T. imaginis* mitochondrial genome (AF335993) as reference contig. The aligned reads were considered for denovo mitochondrial genome of *T. palmi*.

### Genome annotation, visualization, and comparative analysis

The assembled mitogenome was annotated by using MITOS web-server (http://mitos.bioinf.uni-leipzig.de/index.py) to estimate the position of protein coding regions, tRNAs, rRNAs and their secondary structures. The boundaries of PCGs and rRNAs was confirmed manually by BLASTn, BLASTp and ORF Finder in NCBI (https://www.ncbi.nlm.nih.gov/orffinder/). The PCGs were translated into putative proteins using the invertebrate mitochondrial DNA genetic code. The ClustalX program was used to assign the initiation and termination codons in comparison with other thrips reference sequences [19]. MEGAX was used for alignment of the homologous sequences of *T. palmi* with other thrips species [20]. The complete annotated mitogenome of *T. palmi* was prepared using the Sequin submission tool (http://www.ncbi.nlm.nih.gov/Sequin/) for acquiring the accession number from GenBank database. The circular representation of *T. palmi* mitogenome was drawn by CGView online server (http://stothard.afns.ualberta.ca/cgview_server/) with default parameters [21]. The assembled *T. palmi* mitogenome was compared with the other thrips mitogenomes in GenBank database to calculate the nucleotide composition, Relative Synonymous Codon Usage (RSCU), and AT - GC skewness. The nucleotide composition and RSCU were determined using MEGAX [20]. The following formula was used to calculate the skewness: AT skew = (A − T)/ (A + T) and GC skew = (G − C)/(G + C) [22]. The sequence substitution saturation analysis of PCGs was calculated by DAMBE5 software [23]. The secondary structures of tRNAs genes were anticipated by MITOS web server (http://mitos.bioinf.uni-leipzig.de) and confirmed by the tRNAscan-SE (http://lowelab.ucsc.edu/tRNAscan-SE/) [24] and ARWEN 1.2 [25]. The overlapping and intergenic spacer regions of genes in thrips mitogenomes were compared in terms of length and locations. Further the homology of CRs in *T. palmi* and other Thrips mitogenomes were determined through sequence alignment using Clustal Omega [19]. Pairwise analyses of non-synonymous (Ka) and synonymous (Ks) were performed to calculate the selective pressure on PCGs. The nucleotide sequences of each PCG were aligned based on the amino acid sequences in TranslatorX [26] with MAFFT algorithm using GBlocks parameters. The ratios of non-synonymous substitutions (Ka) and synonymous (Ks) substitutions were estimated in DnaSP6.0 [27].

### Phylogenetic analyses

Six complete mitogenomes of five thrips species were retrieved from GenBank on April 1 2018 for comparative study and phylogenetic inference. The complete mitogenome of *Alloeorhychus bakeri* (Hemiptera) was used in the dataset as an outgroup [28]. The phylogenetic relationships among thrips species were estimated with the nucleotide sequences of 13 PCGs. The PCGs were individually aligned in the TranslatorX online platform using the MAFFT algorithm [26] with the GBlocks parameters and default settings. The dataset of all PCGs was concatenated using SequenceMatrix v1.7.845 to form 9696 bp dataset [29]. The optimal substitution model was estimated for the dataset by jModelTest [30]. The dataset was analyzed using maximum likelihood (ML) method implemented in RaxML, and Bayesian inference (BI) method implemented in MrBayes 3.2 [31]. For likelihood analyses, the bootstrap analysis of 1,000 replicates was performed in RaxML [32] with GTR+G+I as best fit model. For BI analyses, two simultaneous runs of 12 million generations were conducted for the dataset using GTR+G+I model and trees were sampled in every 1000 generations, with the first 25% discarded as burn-in. The BI analysis was stopped after reaching the stationary phase and average standard deviation of split frequencies below 0.01. The phylogenetic tree was visualized and edited using FigTree v1.4.2 <u>(http://tree.bio.ed.ac.uk/software/figtree/)</u> [33].

## Results and discussion

### Genome structure, organization and nucleotide composition

The complete mitochondrial genome of *Thrips palmi* was 15,333 base pairs (bp) in length. This is the second largest mitogenome in size among all previously available mitogenomes of insect order Thysanoptera (S1 Table). The mitochondrial genome of *T*. *palmi* was characterized by 37 genes including 13 PCGs, large and small ribosomal RNA (*rrnL* and *rrnS*) genes, 22 transfer RNA (tRNA) genes and two A+T-rich control regions (CRs) (Fig 1). Among all genes, 31 genes were detected on heavy (H) strand and six genes on the light (L) strand (Table 1). The nucleotide composition of complete mitogenome of *T. palmi* was 78.29% AT content (42.71% A + 35.58 % T) and 21.72% GC content (10.14% G + 11.58% C) (Table 2). The AT compositions was observed to be highest 85.81% in control regions (627bp) followed by tRNAs genes 80.26 % (1393 bp), rRNAs genes 80.21% (1870bp) and PCGs 77.43% (11030 bp). Further, the mitogenome of *T. palmi* showed positive AT skewness (0.09) and negative GC skewness (−0.06). The AT skewness of complete mitogenome in other thrips species sequenced so far, ranged from 0.15 (*T*. *imaginis*) to −0.02 (*A*. *obscurus*), while the GC-skewness varies from 0.01 (*F*. *occidentalis*) to −0.11 (*T*. *imaginis*). The positive AT skewness revealed that, the occurrence of Adenine (As) was more than Thiamine (Ts), which has also been observed in other four thrips mitogenomes except *A. obscurus* (−0.02). This positive AT skewness was also observed in rRNA genes, tRNA genes, and control regions of *T. palmi* and were found to be similar with other thrips mitogenomes (Table 2). The negative GC skewness was observed in four thrips species mitogenomes except *F. occidentalis* and *A. obscurus*. The sequence similarity searches of *T. palmi* mitogenome showed the highest similarity (81%) with *T. imaginis* followed by *F. occidentalis* (74%), *F*. *intonsa* (73%), *S. dorsalis* SA1 (72%), *S*. *dorsalis* EA1 (71%) and *A. obscurus* (72%) in GenBank database.

**Fig 1.**
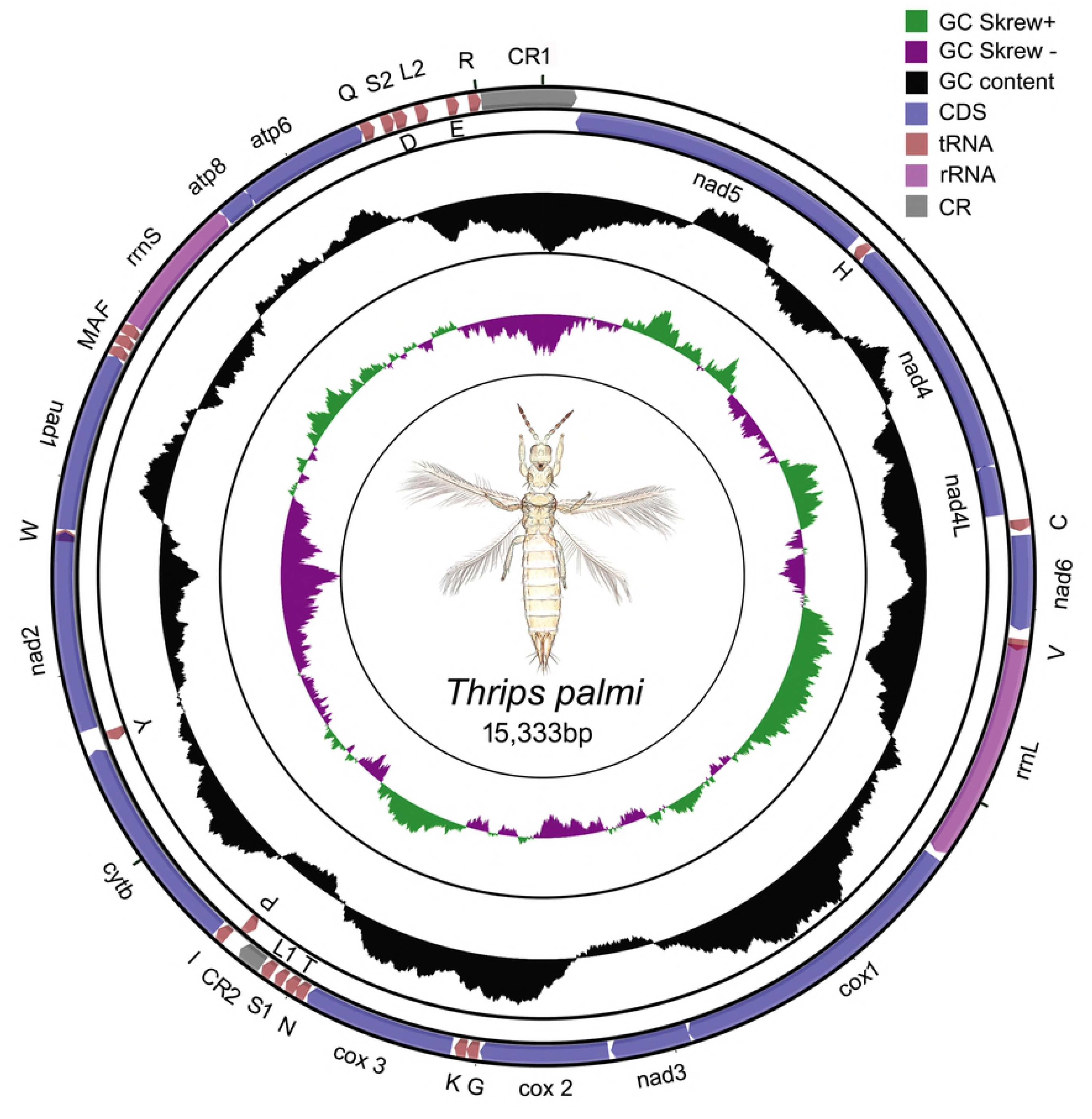
The circular representation of the complete mitochondrial genome of *Thrips palmi*. Direction of gene transcription is indicated by arrows in entire complete genome. PCGs are shown as purple arrows, rRNA genes as pink arrows, tRNA genes as peach color arrows and CR regions as gray rectangles. The GC content is plotted using a black sliding window, as the deviation from the average GC content of the entire sequence. GC-skew is plotted using a colored sliding window (green and orchid color), as the deviation from the average GC skew of the entire sequence. The figure was drawn using CGView online server (http://stothard.afns.ualberta.ca/cgview_server/) with default parameters. The species photograph was taken by second author (KT) using Leica Microscope DM1000 with Leica software application suite (LAS EZ) and edited manually in Adobe Photoshop CS 8.0.

**Table 1.** List of annotated mitochondrial genes of *T. palmi* and its characteristic features. The PCGs and rRNA genes are represented by standard nomenclature, tRNAs are represented as trn followed by the IUPAC-IUB single letter amino acid codes. (+) values in strand represent as heavy (H) and (-) values represent as light (L). IGN represents (+) values as intergenic nucleotides and (-) values as overlapping regions. CR represents the control region.

| Gene | Strand | Location | Size(bp) | Anticodon | Start Codon | Stop Codon | IGN |
| --- | --- | --- | --- | --- | --- | --- | --- |
| *nad5* |  | 176 - 1828 | 1653 | . | ATT | TAA | 29 |
| *trnH* | - | 1858 - 1919 | 62 | CAC | . | . | 2 |
| *nad4* | - | 1922 - 3241 | 1320 | . | ATT | TAA | -7 |
| *nad4l* | - | 3235 - 3510 | 276 | . | ATG | TAG | 34 |
| *trnC* | + | 3545 - 3606 | 62 | UGC | . | . | 19 |
| *nad6* | + | 3626 - 4111 | 486 | . | ATT | TAA | 43 |
| *trnV* | + | 4155 - 4214 | 60 | GUA | . | . | -31 |
| *rrnL* | + | 4184 - 5325 | 1142 | . | . | . | 27 |
| *cox1* | + | 5353 - 6915 | 1563 | . | TTG | TAA | -1 |
| *nad3* | + | 6915 - 7319 | 405 | . | ATG | TAA | 12 |
| *cox2* | + | 7332 - 7988 | 657 | . | ATA | TAA | 5 |
| *trnG* | + | 7994 - 8057 | 64 | GGA | . | . | -1 |
| *trnK* | + | 8057 - 8120 | 64 | AAA | . | . | 13 |
| *cox3* | + | 8134 - 8916 | 783 | . | ATA | TAA | 4 |
| *trnN* | + | 8921 - 8983 | 63 | AAC | . | . | -3 |
| *trnT* | + | 8981 - 9044 | 64 | ACA | . | . | 7 |
| *trnS1* | + | 9052 - 9108 | 57 | AGA | . | . | 15 |
| *trnL1* | + | 9124 - 9188 | 65 | CUA | . | . | 3 |
| CR2 | + | 9192 - 9329 | 138 | . | . | . | 1 |
| *trnP* | - | 9331 - 9394 | 64 | CCA | . | . | 23 |
| *trnI* | + | 9418 - 9481 | 64 | AUC | . | . | 1 |
| *cytb* | + | 9483 - 10595 | 1113 | . | ATA | TAA | 8 |
| *trnY* | - | 10604 - 10666 | 63 | UAC | . | . | 37 |
| *nad2* | + | 10704 - 11726 | 1023 | . | ATA | TAA | -50 |
| *trnW* | + | 11677 - 11738 | 62 | UGA | . | . | 0 |
| *nad1* | + | 11739 - 12662 | 924 | . | ATA | TAA | -4 |
| *trnM* | + | 12659 - 12719 | 61 | AUG | . | . | 1 |
| *trnA* | + | 12721 - 12782 | 62 | GCA | . | . | -1 |
| *trnF* | + | 12782 - 12844 | 63 | UUC | . | . | -2 |
| *rrnS* | + | 12843 - 13570 | 728 | . | . | . | -1 |
| *atp8* | + | 13570 - 13738 | 169 | . | ATA | T(TT) | -7 |
| *atp6* | + | 13732 - 14391 | 660 | . | ATT | TAA | -1 |
| *trnQ* | + | 14391 - 14458 | 68 | CAA | . | . | 41 |
| *trnS2* | + | 14500 - 14564 | 65 | UCA | . | . | 1 |
| *trnD* | + | 14566 - 14633 | 68 | GAC | . | . | 46 |
| *trnL2* | + | 14680 - 14744 | 65 | CUA | . | . | 99 |
| *trnE* | + | 14844 - 14905 | 62 | GAA | . | . | 49 |
| *trnR* | + | 14955 - 15019 | 65 | CGA | . | . | 0 |
| CR1 | + | 15020 -<br>15333+175 | 488 | . | . | . | . |

### Protein coding genes (PCGs) and Relative Synonymous Codon Usage (RSCU)

The mitochondrial genome of *T*. *palmi* was represented by 13 PCGs (*atp6*, *atp8*, *cox1*, *cox2*, *cox3*, *cytb*, *nad1*, *nad2*, *nad3*, *nad4*, *nad4L*, *nad5*, and *nad6*). The ATN initiation codons (six with ATA, four with ATT and two with ATG) was observed in 12 PCGs except *cox1* (TTG). The TTG start codon for *cox1* is unique in *T. palmi* as ATN start codon is observed for *cox1* in all other thrips mitogenomes assembled so far (S2 Table). The PCGs were terminated with TAA stop codon except *nad4L* (TAG) and *atp8* with an incomplete stop codon. The incomplete termination codons are common in other insect mitogenomes which are recovered by the post transcriptional polyadenylation [34–36]. The average AT composition of *T. palmi* PCGs was 77.43% with the highest of 84.06% in *nad4L* gene. However, the average GC composition of the PCGs of *T. palmi* was 22.57%, highest 27.9% in *cox1* gene. The nucleotide composition in all PCGs of *T. plami* showed 41.57% Adenine (A), 35.86% Thiamine (T), 10.09% Guanine (G), and 12.48% Cytosine (C) (Fig 2, S3 Table).

**Fig 2.**
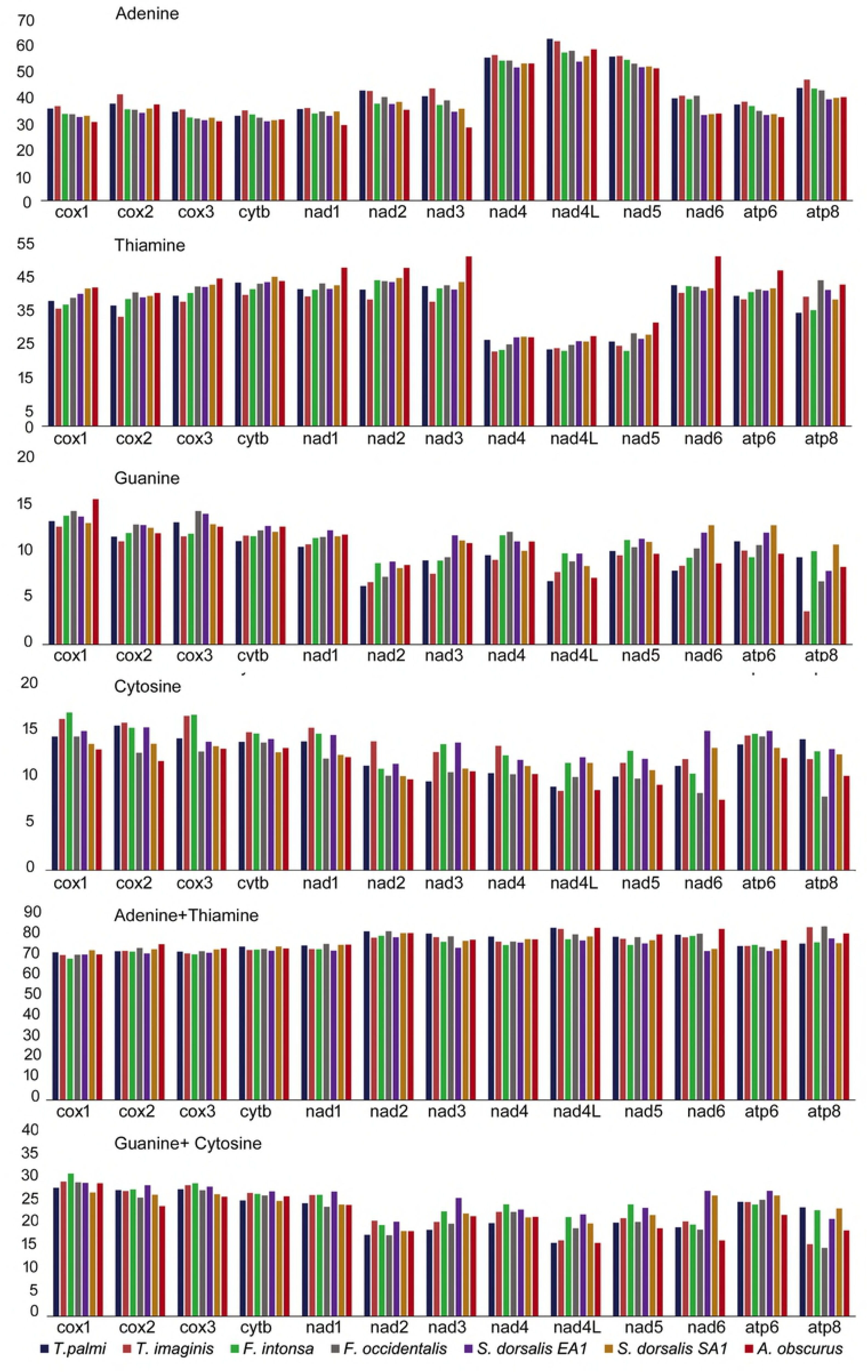
Comparative analysis of nucleotide composition of PCGs within the studied thrips species.

Further, the comparative analysis showed that the adenine (A) composition in all PCGs of seven mitogenomes of six thrips species were varied from 27.68% (*nad3* of *A. obscurus*) to 61.23% (*nad4L* of *T. palmi*), the thiamine (T) composition from 22.23% (*nad4* of *T. imaginis*) to 50.59% (*nad6* of *A. obscurus*), the guanine (G) composition from 3.61% (*atp8* of *T. imaginis*) to15.8% (*cox1* of *A*. *obscurus*), and cytosine (C) composition from 7.65% (*nad6* of *A. obscurus*) to17.10% (*cox1* of *F. intonsa*). The Relative Synonymous Codon Usage (RSCU) data analysis of 3,676 codons in 13 PCGs of *T. palmi* revealed that Phenylalanine (Phe) (15.37%), Leucine (Leu) (14.25%), Isoleucine (Ile) (7.64%), Tyrosine (Tyr) (6.74%), Asparagine (Asn) (6.56%), Lysine (Lys) (5.69%) and Methionine (Met) (4.40%) were the most frequently used amino acids (Fig 3, S4 Table). Comparative RSCU analysis of six thrips species mitogenomes showed that Phe, Leu, Ile, Tyr, Asn, Lys and Met were the seven most frequent amino acids and TTT (Phe), TTA (Leu), ATT (Ile), TAT (Tyr), AAT (Asn) and AAA (Lys), ATA (Met) were the most frequently used codons. In contrast, almost all the frequently used codons ended with A/T, which may lead to the A and T bias in the mitochondrial genomes in thrips species. The codon CGG (Arg) and GCG (Ala) were absent in *T. palmi* and *S. dorsalis* SA1 respectively, while these codons were present in other thrips mitogenomes. Both of the missing codons were preferred to end with ‘G’ in the third codon position (Fig 4). Sequence saturation analysis of PCGs of the thrips mitogenomes exhibited an increasing rate of transitions and transversions ratio along with the divergence value (S1 Fig, S5 Table).

**Fig 3.**
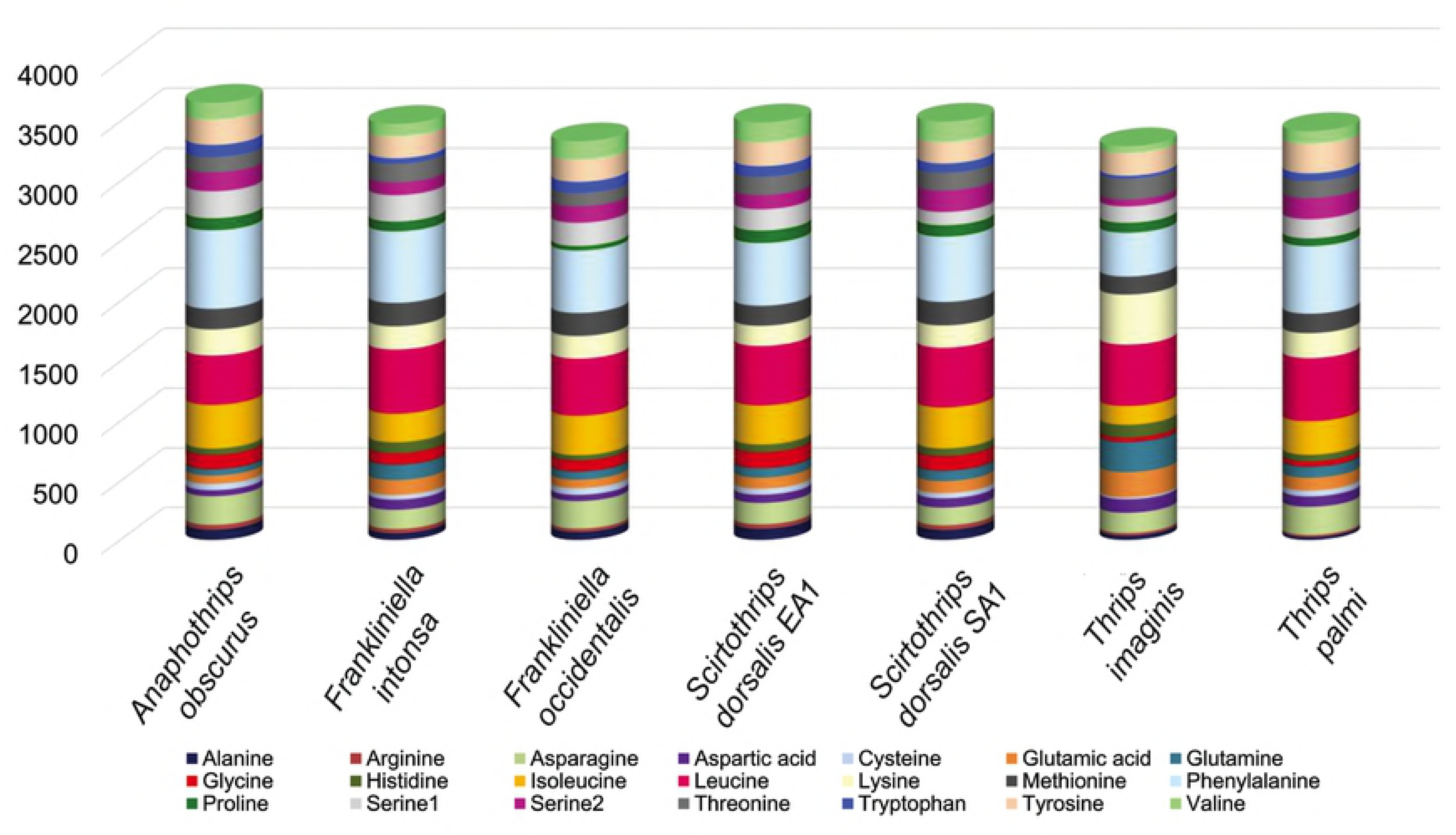
The Relative Synonymous Codon Usage (RSCU) of the mitochondrial genome of *T. palmi* and other thrips species.

**Fig 4.**
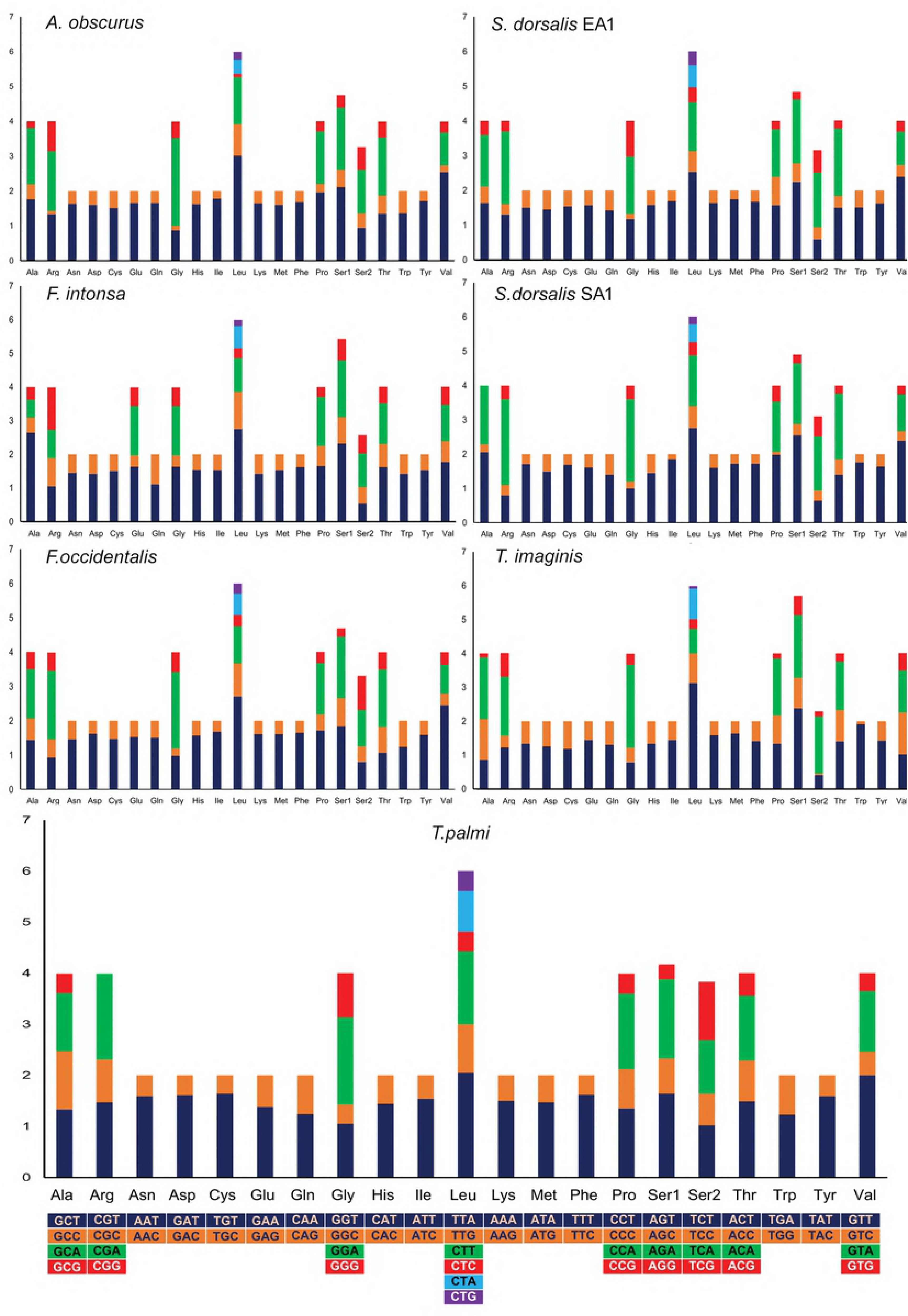
Comparison of codon usage within the mitochondrial genome of members of the thysanopterans.

### Synonymous and non-synonymous substitutions in PCGs

The Synonymous and non-synonymous substitutions (Ka/Ks) ratio is an indicator for investigating the selective pressure and evolutionary relations of the homogenous or heterogeneous species [11]. It was reported that, the Ka/Ks>1 for positive selection, Ka/Ks=1 for neutral mutation, and Ka/Ks<1 for negative selection [37–39]. The Ka/Ks ratio ranges from 0.078±0.02 in *cox1* to 0.913±0.40 in *atp8* gene and the resulted following order: *cox1*<*cox3*<*cytb*<*cox2*<*atp6*<*nad1*<*nad5*<*nad4L*<*nad4*<*nad3*<*nad6*<*nad2*<*atp8*. This result indicated that the 13 PCGs of all thrips mitogenomes including the studied *T. palmi* species were evolving under purifying selection (Fig 5A). Comparative analysis of Ka/Ks ratio among the 13 PCGs of thrips species showed that *atp8* (1.7) and *nad6* (1.3) genes are evolving under positive/relaxed selection with reference to *F. intonsa* and *S. dorsalis* SA1 respectively (Fig 5B). Further, the Ka/Ks was <1 for the remaining PCGs of *T. palmi* with reference to other thrips species, suggested that mutations were replaced by synonymous nucleotides [10]. The lowest Ka/Ks ratio was observed for *cox1* gene representing less variations in amino acids and hence had been evidenced as potential molecular marker for DNA barcoding [8] and phylogeny in thrips [6].

**Fig 5.**
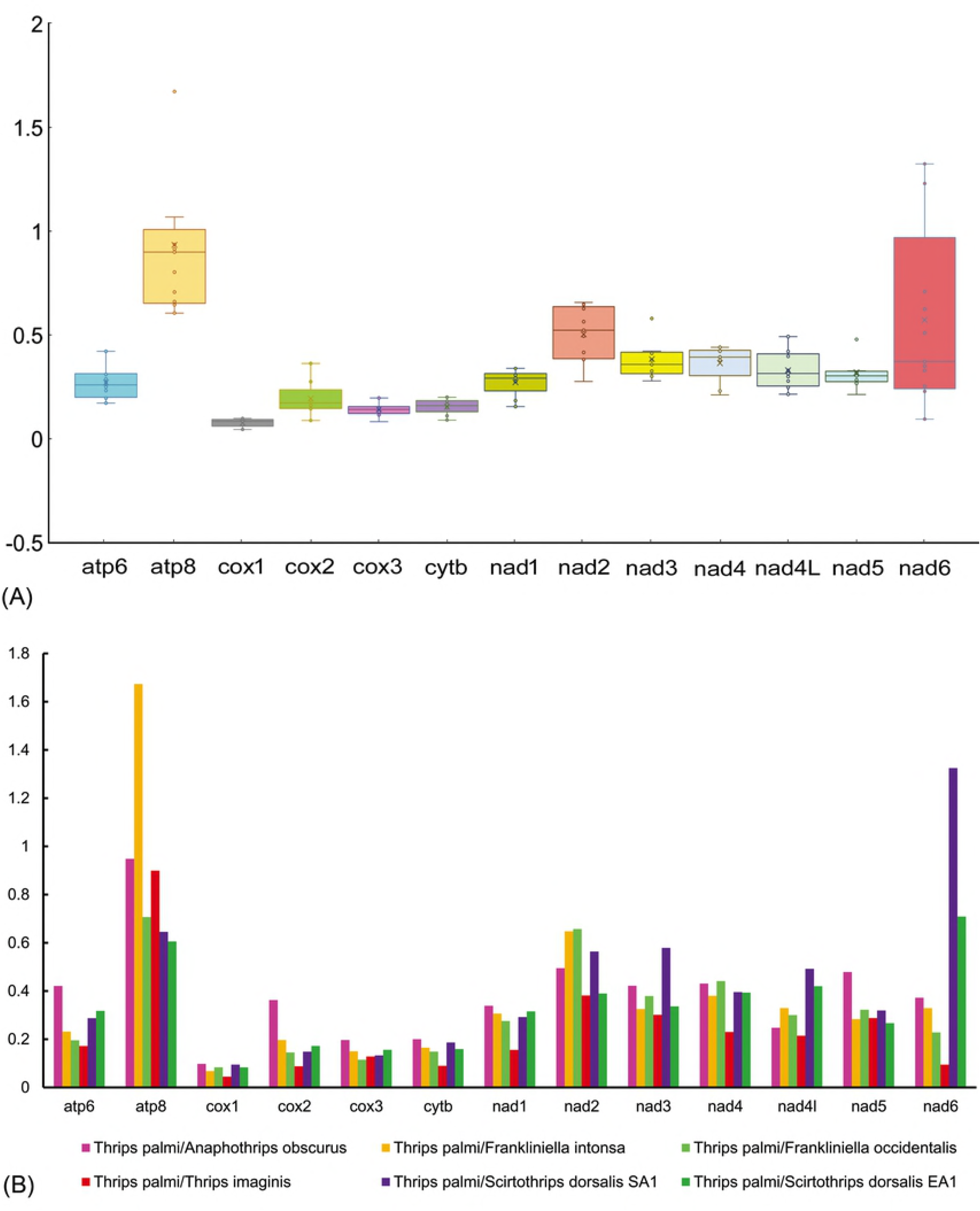
(A) Ratios estimation, box plot for pairwise divergence of Ka/Ks ratio for each one of the mitochondrial PCGs. (B) Evolutionary rates (Ka/Ks) of individuals PCGs of *T. palmi* with other thrips species.

### Ribosomal RNA (rRNA) and transfer RNA (tRNA) genes

The mitogenome of *T. palmi* comprises two rRNA genes as observed in other insect mitogenomes. The large ribosomal gene (rrnL) was 1142 bp long, and located between *trnV* and *cox1*. Further, the small ribosomal gene (rrnS) was 728 bp long, and located between *trnF* and *atp8* gene. The AT content of two rRNA was 80.21% in *T. palmi* which is the highest in comparison with other available thrips mitogenomes. Both AT skewness (0.18) and GC skewness (0.08) of *T. palmi* were positive, that is also similar to other previously sequenced thrips mitogenomes. The locations of rrnL and rrnS were upstream of *cox1* and *atp8* gene, the arrangements seems to be conserved in insect order Thysanoptera. The *T. palmi* mitogenomes contained 22 tRNAs (ranging from 57 to 68 bp in length) with a total length of 1,393 bp. Nineteen tRNA genes were coded by the H-strand and three (*trnY, trnP and trnH*) by the L-strand. The A+T content of tRNAs were 80.26% with positive AT skewness (0.05) and GC skewness (0.03) (Table 2). Most of the tRNA showed the typical cloverleaf secondary structure; absence of DHU stem and loop was observed in *trnV* and *trnS;* absence of DHU loop in *trnE*; lack of TψC loop in *trnN*; variation in TψC arm and loop in other tRNAs (S2 Fig). The absence of DHU stem and loop in *trnV* was consistent in all the thrips species mitogenomes assembled so far. The arrangements of tRNA genes were varied throughout the complete mitogenome of *T. palmi* and other publically available thrips mitogenomes. Among 22 tRNAs genes, 12 genes (*trnA*, *trnL2*, *trnG*, *trnK*, *trnY*, *trnW*, *trnF*, *trnC*, *trnI*, *trnV*, *trnM*, and *trnH*) were found to be conserved within the six thrips species; in their locations respect to the upstream or downstream of PCGs and *rRNA* genes (Fig. 6A). The position of *trnN* seems to be genus specific; between *cox3* and CR2 in the species of genus *Thrips* (*T. palmi* and *T. imaginis*), between *atp6* and CR1 in the species of genus *Frankliniella* (*F. intonsa* and *F. occidentalis*), between *cox3* and *cytb* (*S. dorsalis*) and between *nad3* and *cytb* in *A. obscurus*. The location of *trnS1* (between *cox3* and CR2) is unique in *T. palmi* among all the studied thrips mitogenomes. The position of *trnS2* is observed between *atp6* and CR1 in both *T. palmi* and *T. imaginis*, while it is located in the downstream of *rrnL* in other five thrips mitogenomes. The *trnD* was translocated between *atp6* and CR1 in *T. palmi* and *T. imaginis* from downstream of *cox2* in other five thrips mitogenomes (Fig 6A). The position of *trnR* was found upstream (*A. obscurus*, *F. intonsa*, *F. occidentalis*, *S. dorsalis*) or downstream (*T. imaginis*) of *cox3* gene, however, it was translocated between *atp6* and CR1 in *T. palmi*. The *trnE* was located between *atp6* and CR1 in *T. palmi* along with the *Frankliniella* spp. and *T. imaginis*, while it was found upstream of *cytb* in *S. dorsalis* and *A. obscurus*. The *trnT* was found to be conserved in Thysanoptera mitogenomes except *A. obscurus. The trnL1* in *T. palmi* was found to be similar to *S. dorsalis* (downstream of *cox3*), while it was downstream of *atp6* in *T. imaginis* and *Frankliniella* spp., downstream of *nad4L* in *A. obscurus.* The *trnP* was observed upstream of *cytb* in *Thrips* spp. and *S. dorsalis*; but it was downstream of *cytb* in *Frankliniella* spp. and downstream of *nad6* in *A. obscurus*. The *trnQ* was upstream of *cytb* in *S. dorsalis, F. occidentalis* and *A. obscurus*, but downstream in *F. intonsa*, however, *trnQ* was located downstream of *atp6* in *Thrips* spp. (Fig 6A). The mismatched base pairs (G-U wobble pairs) were observed in seven tRNA genes of *T. palmi* mitogenome. These wobble mismatches were observed in *trnL2* and *trnL1* (in DHU arm), *trnA*, *trnS1*, and *trnL2* (in acceptor arm), *trnS2* and *trnT* (in T ΨC arm).

**Table 2.** Nucleotide Composition and skewness in different Thysanoptera mitogenomes considered for comparative analysis.

| Species | Size(bp) | A% | G% | T% | C% | GC% | AT% | AT skewness | GC skewness |
| --- | --- | --- | --- | --- | --- | --- | --- | --- | --- |
| **Whole mtgenome** |  |  |  |  |  |  |  |  |  |
| *T. palmi* | 15,333 | 42.71 | 10.14 | 35.58 | 11.58 | 21.72 | 78.29 | 0.09 | -0.06 |
| *T. imaginis* | 15,407 | 43.85 | 10.47 | 32.72 | 12.96 | 23.43 | 76.57 | 0.15 | -0.11 |
| *F. intonsa* | 15,215 | 41.24 | 11.06 | 34.68 | 13.01 | 24.07 | 75.93 | 0.09 | -0.08 |
| *F. occidentalis* | 14,889 | 40.98 | 11.35 | 36.62 | 11.06 | 22.41 | 77.60 | 0.06 | 0.01 |
| *S. dorsalis* EA1 | 15,343 | 39.12 | 11.61 | 36.62 | 12.64 | 24.25 | 75.74 | 0.03 | -0.04 |
| *S. dorsalis* SA1 | 15,204 | 39.83 | 11.18 | 37.56 | 11.42 | 22.60 | 77.39 | 0.03 | -0.01 |
| *A. obscurus* | 14,890 | 38.38 | 11.27 | 39.75 | 10.60 | 21.87 | 78.13 | -0.02 | 0.03 |
| **PCG** |  |  |  |  |  |  |  |  |  |
| *T. palmi* | 11030 | 41.57 | 10.09 | 35.86 | 12.48 | 22.57 | 77.43 | 0.07 | -0.11 |
| *T. imaginis* | 10,923 | 42.82 | 9.39 | 34.03 | 13.75 | 23.15 | 76.85 | 0.11 | -0.19 |
| *F. intonsa* | 11009 | 39.70 | 10.90 | 35.65 | 13.77 | 24.66 | 75.34 | 0.05 | -0.12 |
| *F. occidentalis* | 10,852 | 39.78 | 10.99 | 37.81 | 11.41 | 22.40 | 77.60 | 0.03 | -0.02 |
| *S. dorsalis* EA1 | 10,954 | 37.23 | 11.70 | 37.35 | 13.73 | 25.43 | 74.57 | -0.002 | -0.08 |
| *S. dorsalis* SA1 | 10,974 | 38.17 | 11.48 | 38.02 | 12.33 | 23.81 | 76.19 | 0.002 | -0.04 |
| *A. obscurus* | 11168 | 36.95 | 10.81 | 41.26 | 10.98 | 21.78 | 78.22 | -0.06 | -0.01 |
| **tRNA** |  |  |  |  |  |  |  |  |  |
| *T. palmi* | 1,393 | 42.21 | 10.12 | 38.05 | 9.62 | 19.74 | 80.26 | 0.05 | 0.03 |
| *T. imaginis* | 1,540 | 43.72 | 9.54 | 36.72 | 10.02 | 19.56 | 80.44 | 0.09 | -0.02 |
| *F. intonsa* | 1,392 | 43.51 | 10.68 | 35.79 | 10.03 | 20.71 | 79.29 | 0.10 | 0.03 |
| *F. occidentalis* | 1,402 | 42.44 | 10.47 | 37.43 | 9.67 | 20.13 | 79.87 | 0.06 | 0.04 |
| *S. dorsalis* EA1 | 1,426 | 40.69 | 10.93 | 37.52 | 10.86 | 21.79 | 78.21 | 0.04 | 0.00 |
| *S. dorsalis* SA1 | 1,419 | 41.51 | 10.43 | 38.13 | 9.94 | 20.37 | 79.63 | 0.04 | 0.02 |
| *A. obscurus* | 1,430 | 39.87 | 10.69 | 39.80 | 9.63 | 20.33 | 79.67 | 0.00 | 0.05 |
| **rRNA** |  |  |  |  |  |  |  |  |  |
| *T. palmi* | 1,870 | 47.38 | 10.70 | 32.83 | 9.09 | 19.79 | 80.21 | 0.18 | 0.08 |
| *T. imaginis* | 1,876 | 47.65 | 10.77 | 32.14 | 9.43 | 20.20 | 79.80 | 0.19 | 0.07 |
| *F. intonsa* | 1,699 | 47.15 | 11.30 | 32.02 | 9.54 | 20.84 | 79.16 | 0.19 | 0.08 |
| *F. occidentalis* | 1,848 | 45.94 | 12.18 | 33.93 | 7.95 | 20.13 | 79.87 | 0.15 | 0.21 |
| *S. dorsalis* EA1 | 1,775 | 43.21 | 11.89 | 34.99 | 9.92 | 21.80 | 78.20 | 0.11 | 0.09 |
| *S. dorsalis* SA1 | 1,777 | 45.36 | 11.65 | 34.44 | 8.55 | 20.20 | 79.80 | 0.14 | 0.15 |
| *A. obscurus* | 1,812 | 43.16 | 11.70 | 36.59 | 8.55 | 20.25 | 79.75 | 0.08 | 0.16 |
| **Control region** |  |  |  |  |  |  |  |  |  |
| *T. palmi* | 626 | 44.82 | 4.63 | 40.99 | 9.57 | 14.19 | 85.81 | 0.04 | -0.35 |
| *T. imaginis* | 900 | 47.56 | 16.67 | 25.22 | 10.56 | 27.22 | 72.78 | 0.31 | 0.22 |
| *F. intonsa* | 942 | 41.72 | 7.86 | 38.22 | 12.21 | 20.06 | 79.94 | 0.04 | -0.22 |
| *F. occidentalis* | 595 | 40.34 | 7.90 | 43.70 | 8.07 | 15.97 | 84.03 | -0.04 | -0.01 |
| *S. dorsalis* EA1 | 820 | 40.12 | 9.51 | 39.02 | 11.34 | 20.85 | 79.15 | 0.01 | -0.09 |
| *S. dorsalis* SA1 | 767 | 35.33 | 9.26 | 43.55 | 11.86 | 21.12 | 78.88 | -0.10 | -0.12 |
| *A. obscurus* | 145 | 25.52 | 8.97 | 62.76 | 2.76 | 11.72 | 88.28 | -0.42 | 0.53 |

**Fig 6.**
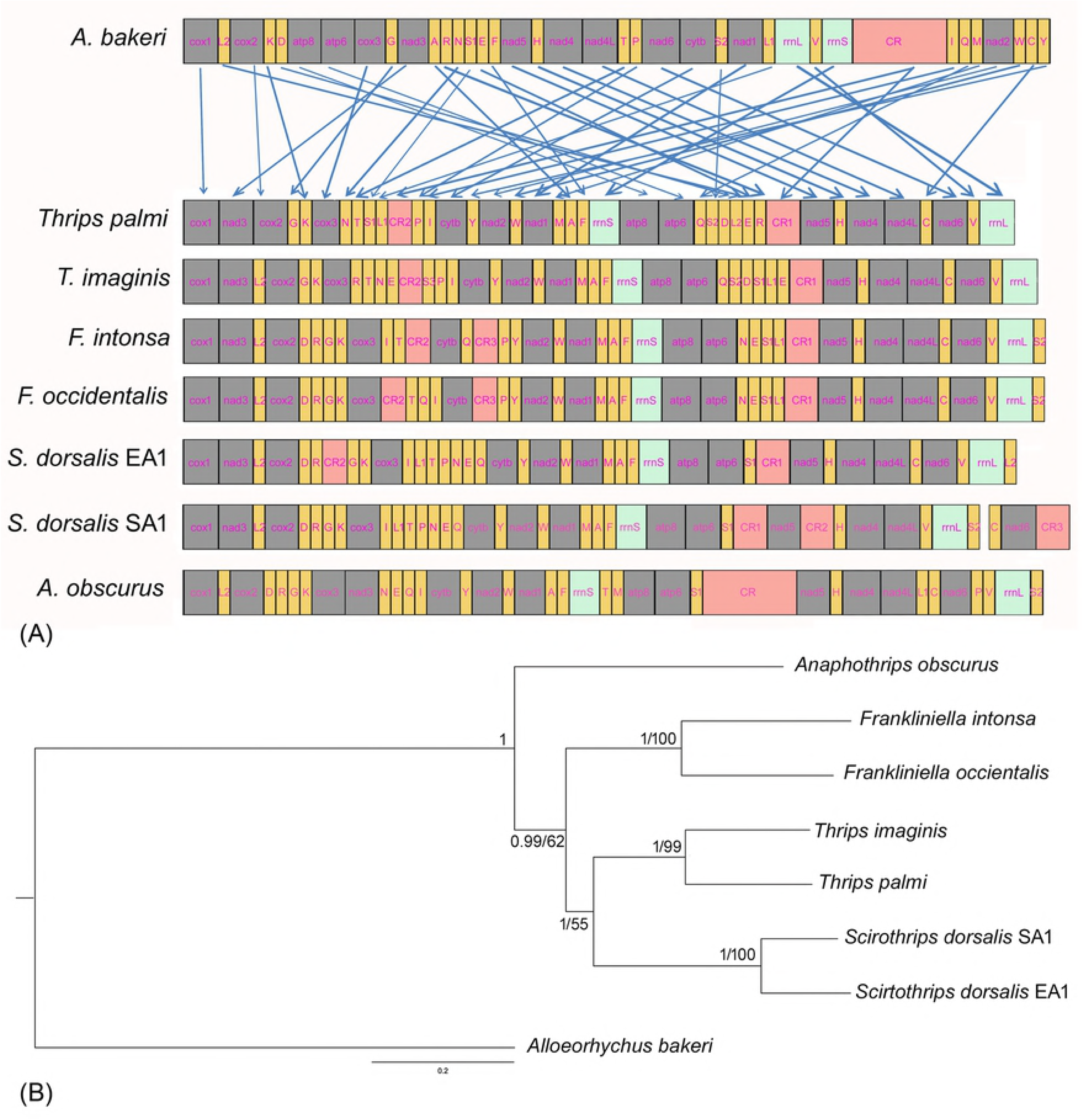
(A) Rearrangements of genes in *T*. *palmi* Mitogenome with respect to other thrips species and ancestor (*A. bakeri*). The PCGs and rRNAs are represented by their standard nomenclature with grey and sea green color respectively. The tRNAs are represented by the IUPAC-IUB single letter amino acid codes and shown in yellow color. The CRs are represented by peach color and light blue arrows indicate the gene translocations. **(B) Bayesian Phylogenetic tree inferred by 13 PCGs of thrips mitogenomes.** The Bayesian posterior probabilities and Maximum likelihood bootstrap supports are superimposed with each node. The tree is drawn to scale with values indicated along with the branches (BI/ML).

### Overlapping and intergenic spacer regions

The mitogenome of *T. palmi* had 24 intergenic spacer regions in a total of 520 bp, varying from 1 to 99 bp in length. There are 14 major intergenic spacers of >10 bp in length were observed (Table 1). The comparative analysis depicted highest intergenic spacer region in *T. palmi* in comparison with other thrips species; 17 intergenic spacer of 217 bp in *T. imaginis*, 14 intergenic spacer of 172 bp in *F. intonsa*, 14 intergenic spacer of 217 bp in *F. occidentalis*, 13 intergenic spacer of 436 bp in *S. dorsalis* EA1, 13 intergenic spacer of 342 bp, 19 intergenic spacer of 309 bp in *A. obscurus*. The longest intergenic spacer (99 bp) was observed between the *trnL2* and *trnE* gene in *T. palmi*. However, the shortest intergenic spacer (1 bp) was observed in four positions: *trnP* and S3, *cytb* and *trnI*, *trnA* and *trnM*, *trnD* and *trnS2*. Further, comparative studies showed that the longest intergenic spacer 150 bp were between trnH and nad4 in *S. dorsalis* EA1. The mitogenome of *T. palmi* contained 12 overlapping regions with a total length of 109 bp. The comparative analysis showed that, the highest (15 overlapping regions of 86 bp) were observed in *S. dorsalis* SA1. The smallest overlapping region (1 bp) was observed in five positions in *T. palmi*: *nad3* and *cox1*, *trnK* and *trnG*, *trnF* and *trnA*, *atp8* and *rrnS*, *trnQ* and *atp6*. The largest overlapping region (50 bp) was observed between *trnW* and *nad2* in *T. palmi*. However, the largest overlapping region (66 bp) were observed between *rrnL* and trnS2 in *F. occidentalis* in comparative studies (S6 Table).

### Control Regions (CRs)

The *T. palmi* mitogenome contained two control regions (CR1 and CR2). The CR1 (488 bp) was lied between *trnR* and *nad5* and CR2 (138 bp) located between the *trnL1* and *trnP* (Fig 6A). Further, the CR2 had 63.77% sequence similarity with CR1, indicating a possible duplication and translocation of control region. The total length of CR is 626 bp, which was higher than *A. obscurus* (145 bp) and *F. occidentalis* (595bp) and lower than other thrips mitogenomes. It is noted that the numbers (1-3) and locations of CRs varied among different thrips species mitogenomes. *T. palmi* had two control regions, similar to *T. imaginis*, and *S. dorsalis* EA1. The location of CR1 upstream of *nad5* gene was suggested to be ancestral condition of the thrips species in subfamily Thripinae [16].

### Phylogenetic analyses

The Maximum likelihood (ML) and Bayesian Inference (BI) phylogenetic trees were constructed based on nucleotide sequences of 13 PCGs. The phylogenetic trees generated using both the methods resulted similar topologies (Fig 6B). The tree clustering revealed that species under genus *Frankliniella* and *Thrips*, were clustered under the respective genus clade. The phylogenetic analyses showed that genus *Thrips* is more closely related to genus *Scirtothrips* as compared to genus *Frankliniella*. The genus *Thrips* and *Frankliniella* forms two large genus groups ‘*Thrips* genus groups’ and ‘*Frankliniella* genus group’ which were supposed to be closely related based on the assumed homology for paired Ctenidia on abdominal segments V-VIII [40]. However, Mound 2002 suggested that these two genus-groups are not closely related based on the chaetotaxy of the abdomen and head [41]. Further, the close relationship between *Scirtothrips* and *Thrips* cannot be supported with their morphology. Till date, mitogenome data on thrips is in its early stages and generation of comprehensive mitogenome data on diverse thrips species from different hierarchical level is needed, to understand the phylogenetic and evolutionary relationships among them.

## Acknowledgement

The authors are thankful to the Director, Zoological Survey of India, Kolkata, for providing necessary facilities, constant support and encouragement throughout the study. The study is financially supported by Zoological Survey of India (ZSI), Kolkata, Ministry of Environment Forest and Climate Change (MoEF&CC), New Delhi under National Faunal Genome Resources (NFGR) Program. This work is a part of the Ph. D thesis of the first author (RC).

## Supporting information

**S1 Fig. Transition (S) and transversion (V) saturation plots for PCGs dataset of *T. Palmi*.**

**S2 Fig. Putative secondary structures of the 22 tRNA genes of *T. palmi* mitogenome.** The tRNAs are represented by full names and IUPAC-IUB single letter amino acid codes. The details of stem and loop is mentioned for one tRNA Serine which is applicable for all tRNAs secondary structures.

**S1 Table. Details of the Thysanoptera mitogenomes generated till date and considered**

**for comparative mitogenome study.**

**S2 Table. Start and Stop codons of the PCGs of the thrips mitgenomes.**

**S3 Table. Nucleotide composition in different mitochondrial locus of the studied thrips mitogenomes.**

**S4 Table. RSCU analysis of the PCGs of in *T. palmi* mitogenome.**

**S5 Table. Genetic distance versus transition-transversion ratio of 13 PCGs in *T. palmi* mitogenome.**

**S6 Table. Comparison of intergenic nucleotides (INs) and overlapping regions of each gene in thrips mitogenomes.**

